# Hematopoietic-specific heterozygous loss of *Dnmt3a* exacerbates colitis-associated colon cancer

**DOI:** 10.1101/2022.12.30.522355

**Authors:** Yang Feng, Rachel C Newsome, Troy Robinson, Robert L Bowman, Ashley N Zuniga, Kendra N Hall, Cassandra M Bernsten, Daniil E Shabashvili, Kathryn I Krajcik, Chamara Gunaratne, Zachary J Zaroogian, Kartika Venugopal, Heidi L Casellas Roman, Ross L Levine, Walid K Chatila, Rona Yaeger, Alberto Riva, Daniel Kopinke, Christian Jobin, Dorina Avram, Olga A Guryanova

## Abstract

Clonal hematopoiesis (CH) is defined as clonal expansion of mutant hematopoietic stem cells absent diagnosis of a hematologic malignancy. Presence of CH in solid tumor patients, including colon cancer, correlates with shorter survival. We hypothesized that bone marrow-derived cells with heterozygous loss-of-function mutations of *DNMT3A*, the most common genetic alteration in CH, contribute to the pathogenesis of colon cancer.

In a mouse model that combines colitis-associated colon cancer with experimental CH driven by *Dnmt3a^+/Δ^*, we found higher tumor penetrance and increased tumor burden compared to controls. Histopathological analysis revealed accentuated colonic epithelium injury, dysplasia and adenocarcinoma formation. Transcriptome profiling of colon tumors identified enrichment of gene signatures associated with carcinogenesis, including angiogenesis. Treatment with the angiogenesis inhibitor axitinib eliminated the colon tumor-promoting effect of experimental CH driven by *Dnmt3a* haploinsufficiency. This study provides conceptually novel insights into non-tumor-cell-autonomous effect of hematopoietic alterations on colon carcinogenesis and identifies potential therapeutic strategies.

**SUMMARY:** A pre-clinical mouse model demonstrates that genetic alterations in the blood system characteristic of clonal hematopoiesis (CH) contribute to an aggressive solid tumor phenotype. It further identifies cancer angiogenesis as a potential therapeutic target to mitigate adverse CH effects.

## INTRODUCTION

Clonal hematopoiesis (CH) is defined as expanded hematopoietic clone in the absence of an overt hematologic malignancy (1–4). CH is most frequent in the elderly (10-40%), and is commonly driven by somatic mutations in leukemia-associated genes such as *DNMT3A* (5–14). While it seems intuitive that CH is associated with an increased risk of leukemia (8,15–21), growing evidence shows that it is also linked to multiple disease conditions outside of the hematopoietic system, and increased overall mortality (9,22–26). This includes cardiovascular disease (CVD) (23,27–29), infections (30), ulcerative colitis (31), and a wide spectrum of solid tumors (27,32–39). In patients with solid tumors, the prevalence of CH similarly increases with age yet is notably more frequent (~30% of cases) and correlates with shorter survival due primarily to solid tumor progression (1,34,40). The effect is the strongest for CH with presumed leukemic driver mutations such as in the epigenetic modifier gene *DNMT3A*, and increases with CH clone size.

DNMT3A, a *de novo* DNA methyltransferase that epigenetically enforces hematopoietic stem cell differentiation programs (41–47), is recurrently mutated in hematologic malignancies (48–55). *DNMT3A* is by far the most frequently altered gene in CH (8–11,34,56,57) with the majority of mutations consistent with a heterozygous loss of function (LOF, ~50%, truncating indels, splice, and nonsense) (48,52). Yet, despite important clinical implications, the causal relationship between presence of CH and aggressive phenotype of unrelated solid tumors has not been rigorously addressed.

Colon cancer is one of the leading causes of cancer-related deaths in developed countries (58–60). Inflammatory bowel disease (IBD), including ulcerative colitis and Crohn’s disease, is a well-known risk factor for colon cancer (61). Colitis-associated colon cancer (CAC) accounts for 15% of overall mortality among all IBD patients. Compared to sporadic colorectal cancer, patients with CAC often present with multifocal tumors arising from pre-cancerous lesions that are challenging to detect and remove endoscopically and tend to rapidly develop chemoresistance (62). Over 20% of patients with colon cancer in the publicly available Memorial Sloan Kettering Cancer Center clinical sequencing database have detectable CH (63,64), which is notably more prevalent than in age-matched cancer-free population. Despite screening and lifestyle interventions, most patients present with advanced disease associated with poor outcome. To inform the choice of optimal therapeutic approaches and improve survival rates, better understanding of disease-modifying factors that contribute to the aggressive tumor phenotype is critically needed.

Potential involvement of CH in the pathogenesis of coincident solid tumors has far-reaching translational impact, yet better understanding of this relationship is hindered by the lack or animal models. Here, we combined a well-established induction of CAC (65–67) with a bone marrow transplantation (BMT) approach to experimental CH driven by heterozygous *Dnmt3a* loss (42,68) (**Figure 1A,B**). This unique tool enables rigorous interrogation into the role of CH in the pathogenesis of solid tumors driven by unrelated genetic alterations, uncoupled from environmental, lifestyle, iatrogenic, or other confounding factors (34,43,69–72). While attempts to unveil the role for CH in non-malignant conditions such as atherosclerosis or gout were made (23,26,73–75), similar studies in cancer models are limited. In this proof-of-concept study, we report that heterozygous inactivation of *Dnmt3a* restricted to the hematopoietic compartment exacerbates the development of colon cancer phenotype in an inflammation-induced mouse model. We further credential an anti-angiogenesis drug axitinib as a potential therapeutic mitigation strategy.

**Figure 1.**
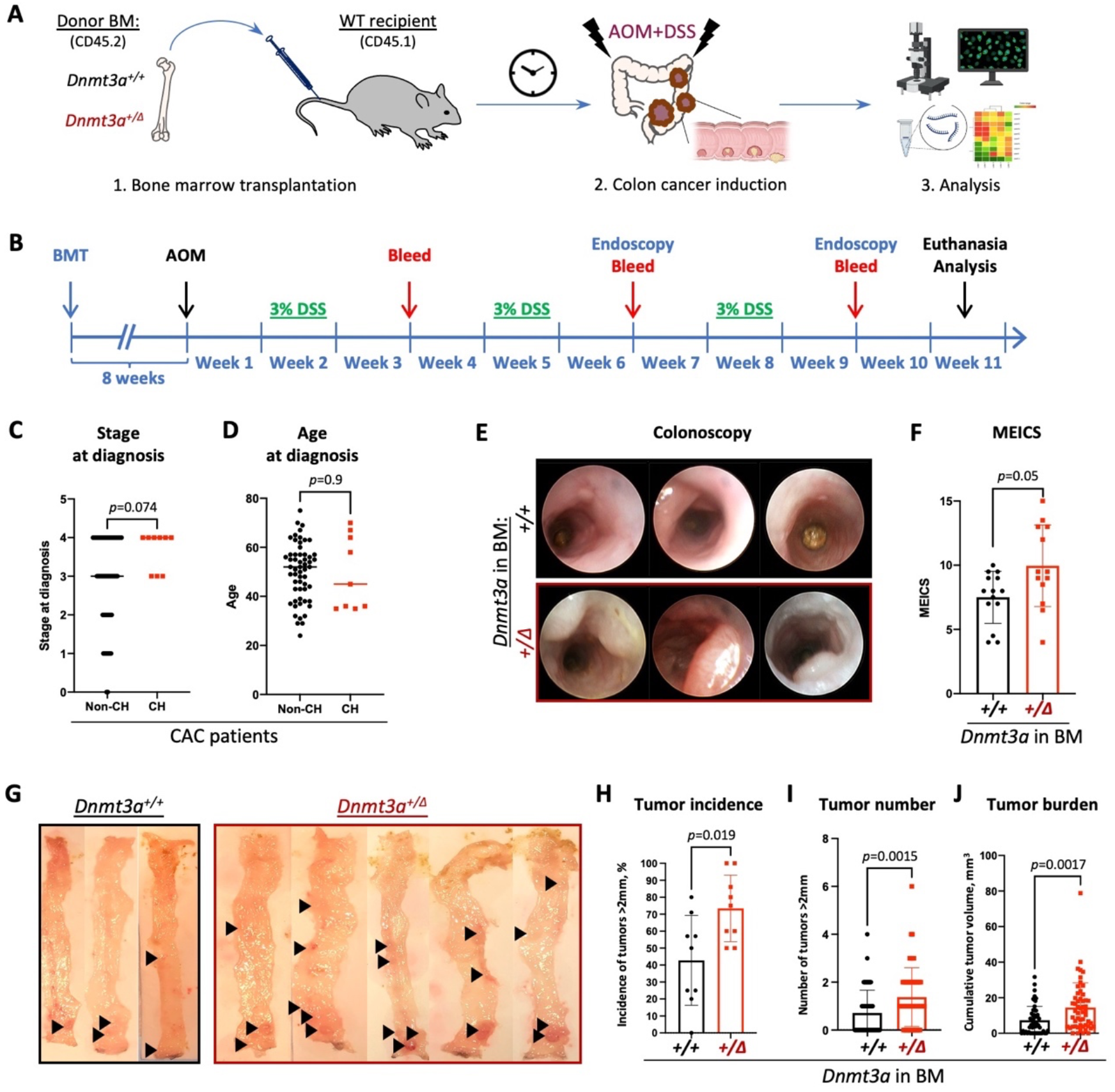
*Dnmt3a* haploinsufficiency specifically in the blood system promotes tumorigenesis in a model of colitis-associated colon cancer. (**A, B**) Experimental workflow using a mouse model of BMT-based CH and AOM/DSS induced CAC (**A**) and timeline (**B**). (**C**) CAC patients with detectable CH at diagnosis (n=9) tend to be more likely to present with an advanced stage of the disease (stage 3 or 4) than those without CH (n=57) (Mann-Whitney test, p=0.074); (**D**) Age of CAC patients with or without CH at diagnosis (n=57 or 9, Mann-Whitney test, p=0.9). (**E, F**) Representative colonoscopy findings by CoIoview mini-endoscopic system in a mouse model of CH (**E**) and corresponding murine endoscopic index of CAC severity (MEICS) scores in *Dnmt3a*^+/+^ and *Dnmt3a^+/Δ^* BM chimerae (**F**, n=14&13, Student’s *t* test, p=0.05). (**G-J**) Representative gross pathology of colons with tumors (black arrowheads, **G**), proportion of animals presenting with tumors (>2mm in diameter) in 9 independent experiments (**H**, Mann-Whitney test, p=0.019), number of large tumors (>2mm in diameter) per animal (**I**, n=55&53, Mann-Whitney test, p=0.0015), and cumulative tumor burden per animal (**J**, n=55&53, Mann-Whitney test, *p*=0.0017).

## RESULTS

### Heterozygous loss of *Dnmt3a* in the bone marrow leads to accentuated CAC phenotype

To explore the relationship between genetic alterations in the blood system and the severity of coincident colitis-associated colon cancer (CAC) we examined presence of CH mutations in paired blood and tumor samples in a cohort of 66 patients with CAC treated at Memorial Sloan-Kettering Cancer Center. We observed a strong trend towards a more advanced disease at diagnosis among patients with detectable CH by MSKCC-IMPACT (63,64) (**Figure 1C**, *p*=0.057, two-sided Chi-square test comparing patients with local (stages 0-1-2) and advanced (stages 3-4) disease, and **Table 1**). Patient age at diagnosis could not explain the difference in CH prevalence (**Figure 1D**), consistent with previous reports implicating that the impact of CH on solid tumor outcomes was not due to differences in age. To directly test the causal relationship between presence of CH and CAC pathogenesis, we performed a proof-of-concept animal study.

**Table 1.**
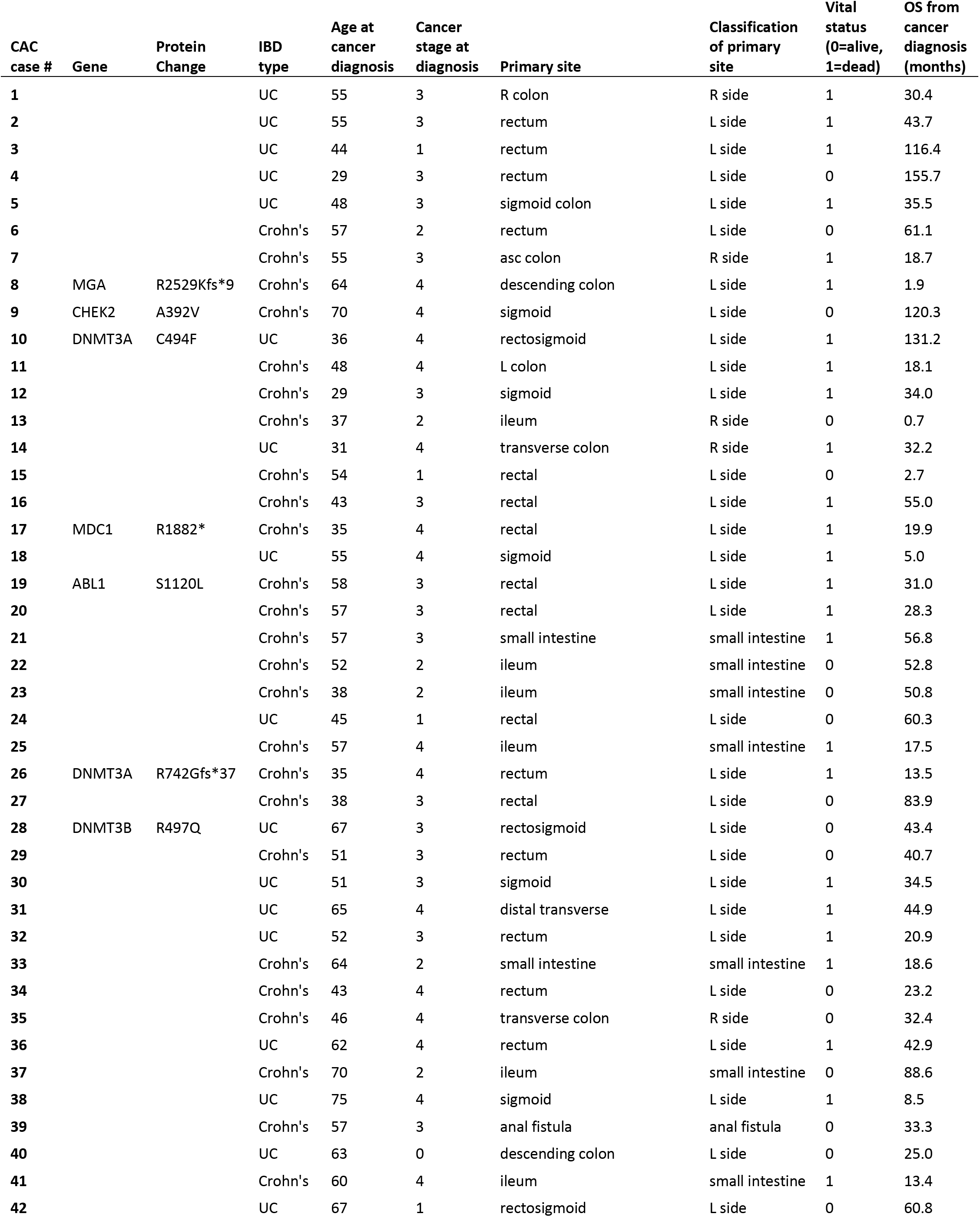
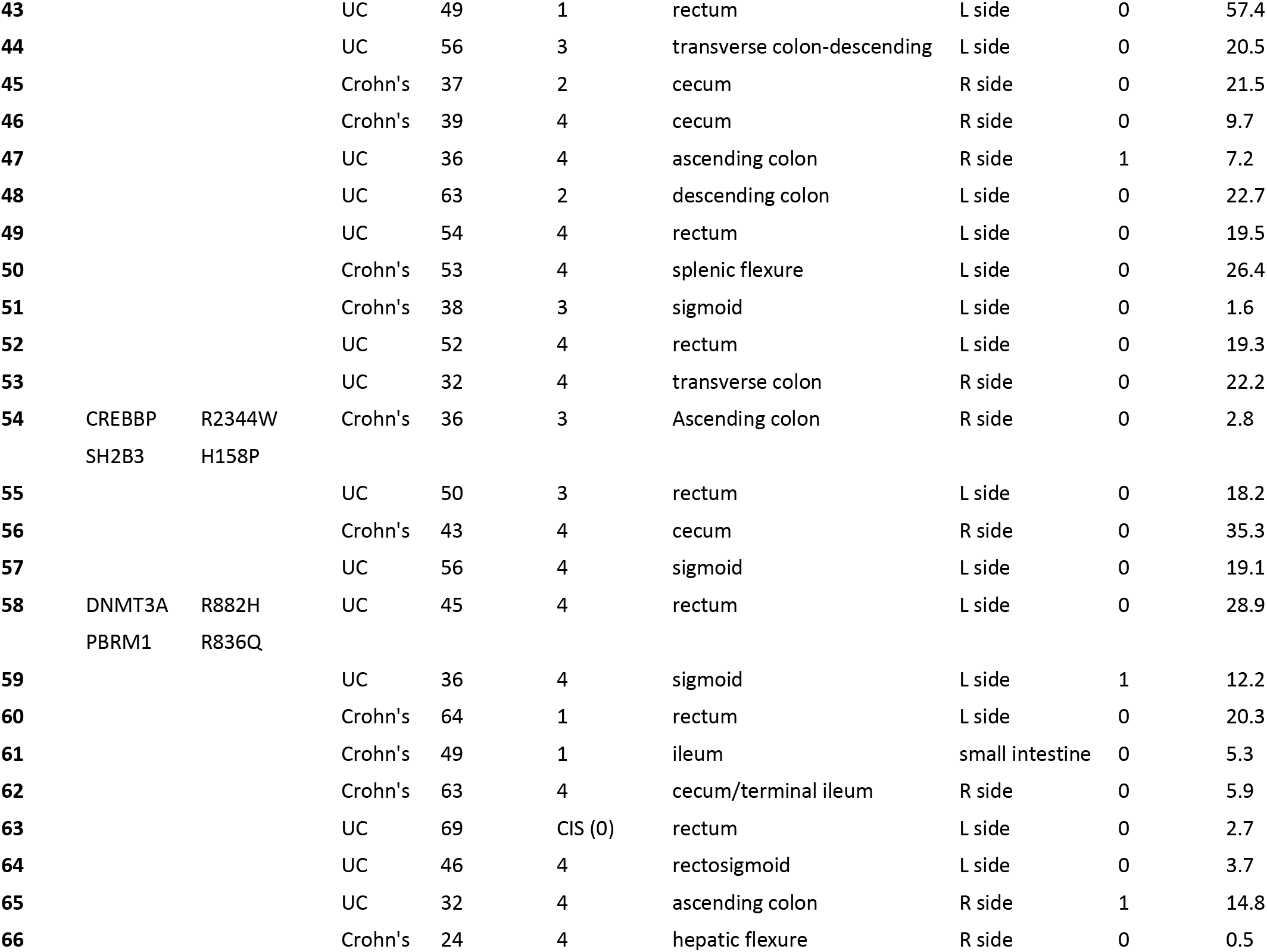
Clinical characteristics and presence of CH in a cohort of 66 patients with CAC.

To experimentally model CH in mice, we adopted a BMT-based approach using *Dnmt3a^+/f^:Mx1-Cre^+^* and *Dnmt3a^+/+^:Mx1-Cre^+^* (WT control) mice as donors after successful Cre-recombination induced by intraperitoneal (IP) administration of poly(I:C), yielding *Dnmt3a^+/Δ^* and *Dnmt3a*^+/+^ mice. Fully engrafted *Dnmt3a^+/Δ^* and control *Dnmt3a*^+/+^ BM chimerae underwent a CAC induction protocol initiated by a single injection of AOM (10mg/kg, IP) followed by three cycles of DSS exposure (3% in sterile drinking water) (76,77) (**Figure 1A,B**). Colonoscopy at 9 weeks after initiation of the AOM/DSS treatment demonstrated increased colon wall opacity, visible bleeding, numerous fibrin patches indicating heightened colon pathology, and detected more, larger tumors in accessible colon regions in *Dnmt3a^+/Δ^* BM chimerae compared to *Dnmt3a*^+/+^-engrafted controls (**Figure 1E**). Modified murine endoscopic index of CAC severity (MEICS) (78) independently scored by two blinded investigators was significantly elevated in the *Dnmt3a^+/Δ^* group (**Figure 1F**).

To further investigate the effect of *Dnmt3a* heterozygous loss in the hematopoietic system characteristic of CH on CAC induction, mice were euthanized for comprehensive colon evaluation 10 weeks after initiation of the AOM and DSS treatment. Colon gross pathology (**Figure 1G**) demonstrated significantly elevated penetrance of the tumor phenotype (incidence of large tumors >2mm in diameter, **Figure 1H**) and higher large tumor number (**Figure 1I**) and increased total tumor burden per colon (**Figure 1J**) in *Dnmt3a^+/Δ^* chimerae. Together, these findings suggest that hematopoietic-specific *Dnmt3a* haploinsufficiency promotes both cancer initiation and progression in the context of CAC.

### *Dnmt3a* haploinsufficiency in the bone marrow leads to accentuated pathological features of CAC

Given that *Dnmt3a^+/Δ^* chimerae have higher tumor burden and larger tumor size compared to wild-type controls, we performed histopathology analysis on colons from both groups based on modified quantitative scoring system for DSS-induced murine CAC (79–82). H&E-stained paraffin sections of swiss-rolled colons showed marked immune infiltration, extensive ulceration and dysplasia of colonic epithelium, and more frequent adenocarcinoma formation with occasional submucosal invasion in *Dnmt3a^+/Δ^*-reconstituted animals (**Figure 2A**). In comparison, wild-type control chimerae showed moderate dysplasia, fewer adenoma polyps, mild ulceration, and more tissue regeneration. Histology scores based on four parameters (immune infiltration (0-3), ulceration (0-3), morphology of colonic epithelium (0-4), and neoplasms (0-4)) independently assigned by two blinded investigators were significantly higher in the *Dnmt3a^+/Δ^* group than in WT (**Figure 2B,C**). We found that in *Dnmt3a^+/Δ^* BM chimerae, increased proliferation of colonic epithelium occurred early in the carcinogenesis process, evidenced by a higher proportion of Ki67-marked cells per crypt in the mucosal layer after the first cycle of DSS (**Figure 2D,E**). Overall, *Dnmt3a* haploinsufficiency in the BM yields more advanced CAC histopathology, consistent with a more severe tumor burden.

**Figure 2.**
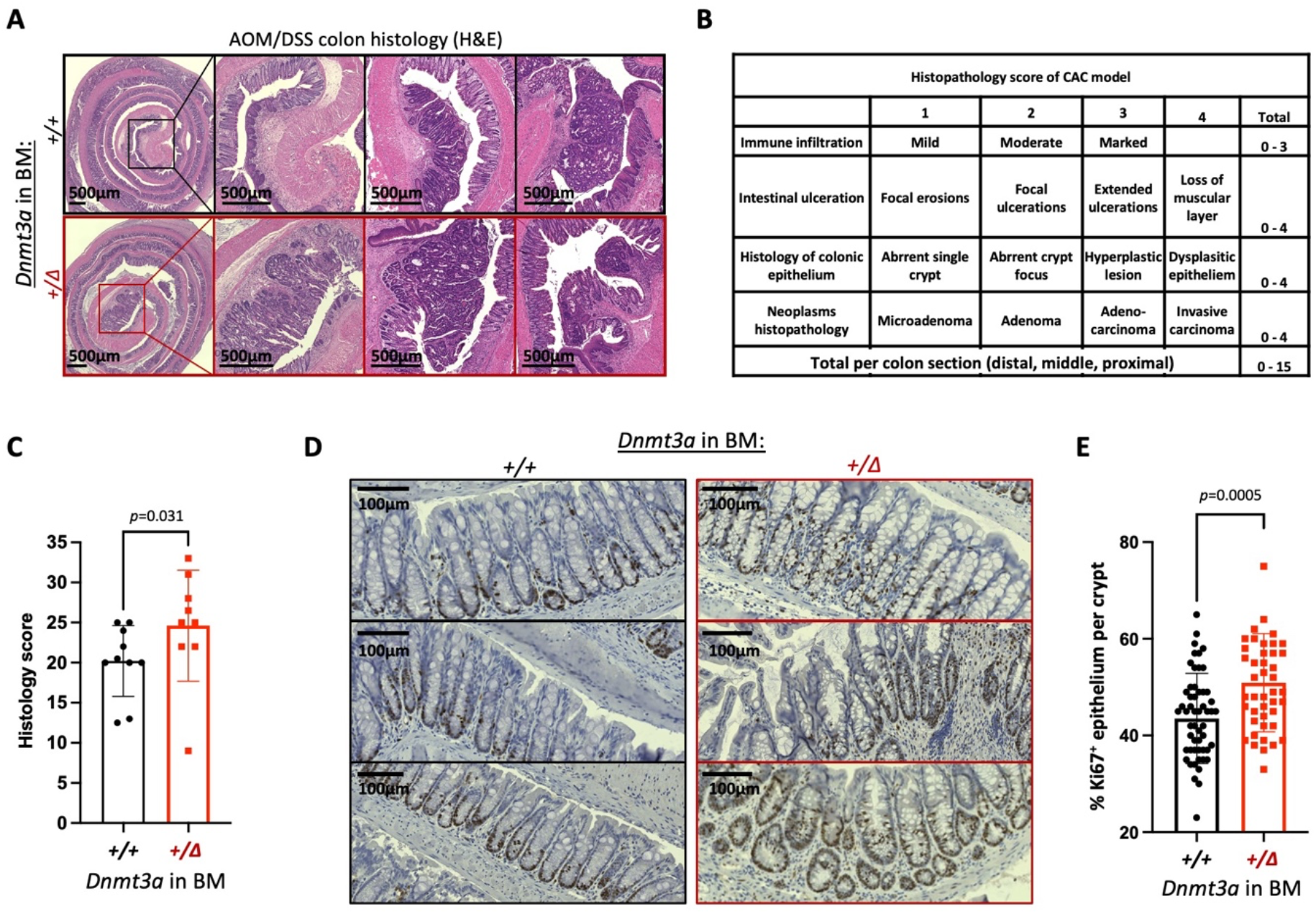
Histopathological features of a more aggressive CAC in animals with hematopoietic-specific heterozygous *Dnmt3a* loss. (**A**) Representative images of H&E-stained swiss-rolled colons. Colons from animals engrafted with *Dnmt3a^+/Δ^* BM exhibit marked immune cell infiltration, dysplastic epithelium, extensive ulceration, global hyperplasia and occasional invasive adenocarcinoma. Bar – 500μm. (**B**) Histopathology scoring criteria. (**C**) Overall histology scores indicate more advanced colon and tumor pathology in *Dnmt3a^+/Δ^* BM-chimerae compared to WT-grafted controls (n=10&9, Mann-Whitney test, *p*=0.031). (**D, E**) Representative examples of Ki67 immunohistochemistry staining in intestinal crypts after one cycle of DSS treatment (**D**, bar – 100μm) and increased proliferation of colonic epithelium in animals with *Dnmt3a^+/Δ^* BM (right panels), quantified as percentage of Ki67* epithelial cells per crypt (**E**, n=55&44, Mann-Whitney test, *p*=0.0005).

### Gene expression profiling identifies signatures of accentuated CAC tumorigenesis in animals with experimental CH driven by *Dnmt3a^+/Δ^*

To identify specific molecular mechanisms likely driving accentuated CAC tumorigenesis in animals with *Dnmt3a^+/Δ^* hematopoiesis, we profiled tumor transcriptomes from *WT* and *Dnmt3a^+/Δ^* chimerae by bulk RNA-seq (**Supplementary Table S1**). We identified 297 differentially expressed genes (log_2_ fold change (log2FC) >1, *p*adj<0.05, **Figure 3A**) with most being upregulated rather than downregulated in the *Dnmt3a^+/Δ^* group (241 up and 56 down, **Supplementary Table S2**). Gene set enrichment analysis (GSEA) (83) using the HALLMARK collection of gene signatures (84) (**Figure 3B**) detected significant positive enrichment of cancer- and proliferation-related pathways such as epithelial-mesenchymal transition, Wnt/β-catenin, and angiogenesis (**Figure 3C-E**), along with E2F, mitotic spindle, G2/M checkpoint, and MYC signaling, and shifts in metabolism such as negative enrichment of oxidative phosphorylation (**Figure 3F**) and adipogenesis/fatty acid metabolism (**Supplementary Table S3**). Elevated expression of the Wnt/Δ-catenin pathway genes is notable (**Figure 3C**) given its known role in driving colon cancer and association with poor prognosis (85). These results are consistent with our findings of increased proliferation within the epithelial layer of regenerating colons in *Dnmt3a^+/Δ^* chimeric mice likely leading to a more advanced CAC pathology.

**Figure 3.**
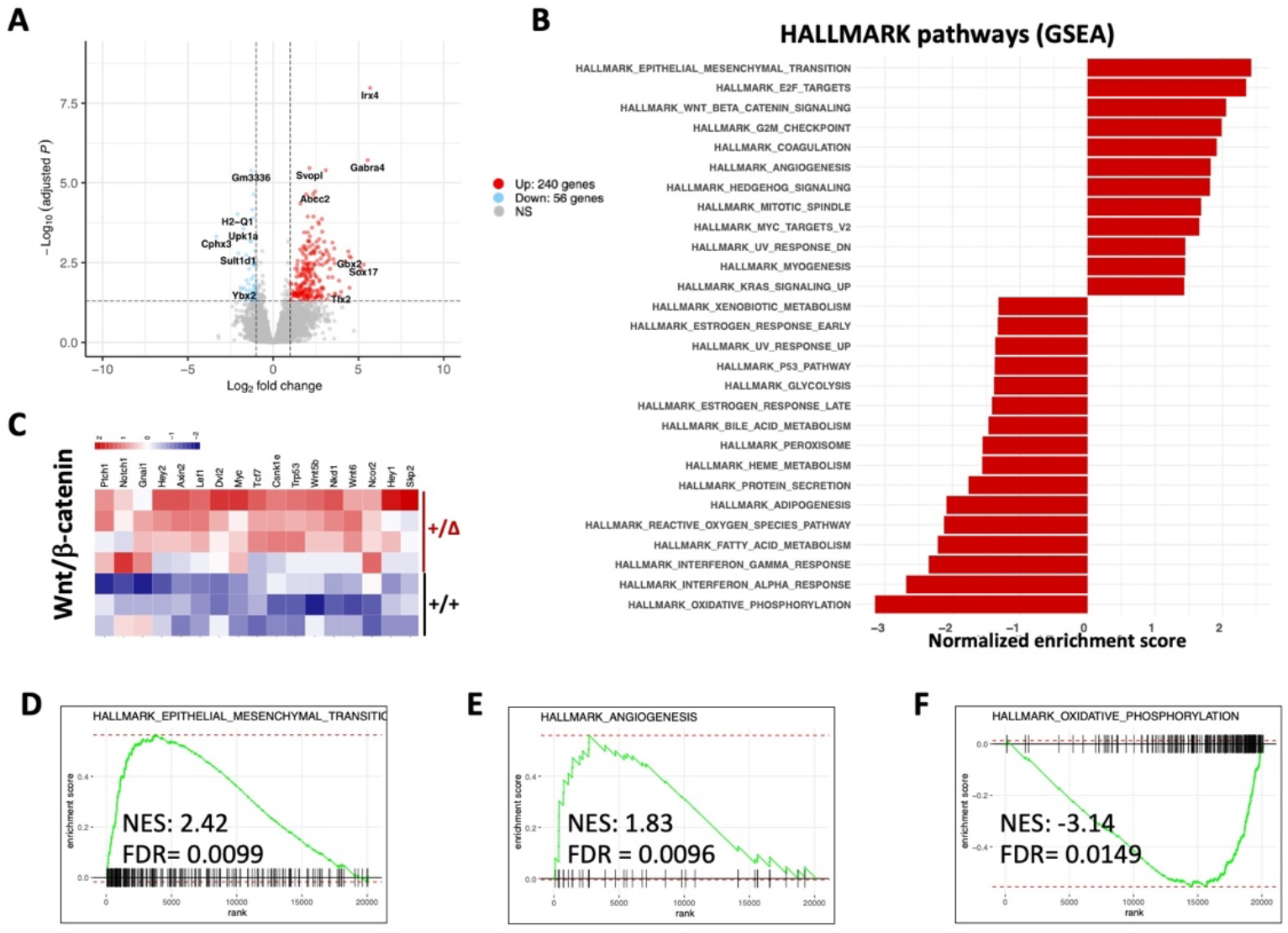
Transcriptomic analysis identifies enrichment of gene expression signatures associated with accentuated colon tumorigenesis, angiogenesis, and changes in metabolism. (**A**) Differential gene expression in colon tumors from animals with and without experimental *Dnmt3a^+/Δ^* CH. Volcano plot showing 297 significantly (| log_2_ fold change |>1, adjusted *p*<0.05) up- (241) or down-regulated (56) genes in *Dnmt3a^+/Δ^* chimerae (n=4) compared to WT control tumors (n=3). (**B**) Most significantly enriched gene sets from the HALLMARK collection (MSigDB) in tumor transcriptomes from *Dnmt3a^+/Δ^* chimerae compared to control *Dnmt3a^WT^*-engrafted mice, | NES|>1.3. (**C**) Heat map of differentially expressed genes showing activation of the Wnt/Δ-catenin pathway in colon tumors from mice with *Dnmt3a^ł/Δ^* bone marrow. (**D-F**) GSEA plots showing positive enrichment of the EMT (**D**) and angiogenesis (**E**) gene signatures and negative enrichment of the oxidative phosphorylation pathway (**F**).

### Treatment with an angiogenesis inhibitor eliminates the tumor-promoting effect of *Dnmt3a^+/Δ^* BM

To further investigate our finding of an enriched angiogenesis-related gene expression signature detected by RNA sequencing, we performed immunofluorescent staining of colon tumors using anti-CD31 antibody that marks endothelia. CAC tumors from animals grafted with *Dnmt3a^+/Δ^* BM had increased CD31 staining compared to *Dnmt3a*^+/+^-reconstituted controls, consistent with more extensive vascularization (**Figure 4A**, red) known to promote tumor growth. Hence, we hypothesized that cancer angiogenesis in animals with experimental CH may be targetable therapeutically to mitigate accentuated tumor phenotype. Axitinib is an orally bioavailable small molecule angiogenesis inhibitor which is FDA approved for treating advanced renal carcinoma with clinical trials for other cancers ongoing (86). Axitinib is a tyrosine kinase inhibitor of vascular endothelial growth factor (VEGF) receptors −1, −2, and −3 that is also active against platelet-derived growth factor receptor (PDGFRβ) and stem cell factor receptor c-KIT/CD117 as part of its potent anti-angiogenic mechanism of action (87). Here, we tested if axitinib abolishes the tumor-promoting effect of *Dnmt3a^+/Δ^* BM during CAC induction (**Figure 4B**). After 10 weeks of axitinib treatment, a significantly lower proportion of *Dnmt3a^+/Δ^*-BM grafted animals had large tumors compared to vehicle controls (**Figure 4C**). Treated animals in the *Dnmt3a^+/Δ^*-BM group developed dramatically fewer large tumors and presented with significantly lower tumor burden per colon (**Figure 4D,E**), while the tumor phenotype in the *Dnmt3a*^+/+^-grafted mice was only slightly affected by axitinib administration. Thus, treatment with an anti-angiogenesis drug axitinib reversed the heightened CAC tumor phenotype in *Dnmt3a^+/Δ^*-chimerae, suggesting its potential translational utility for mitigating unfavorable effects of CH in cancer patients.

**Fig. 4.**
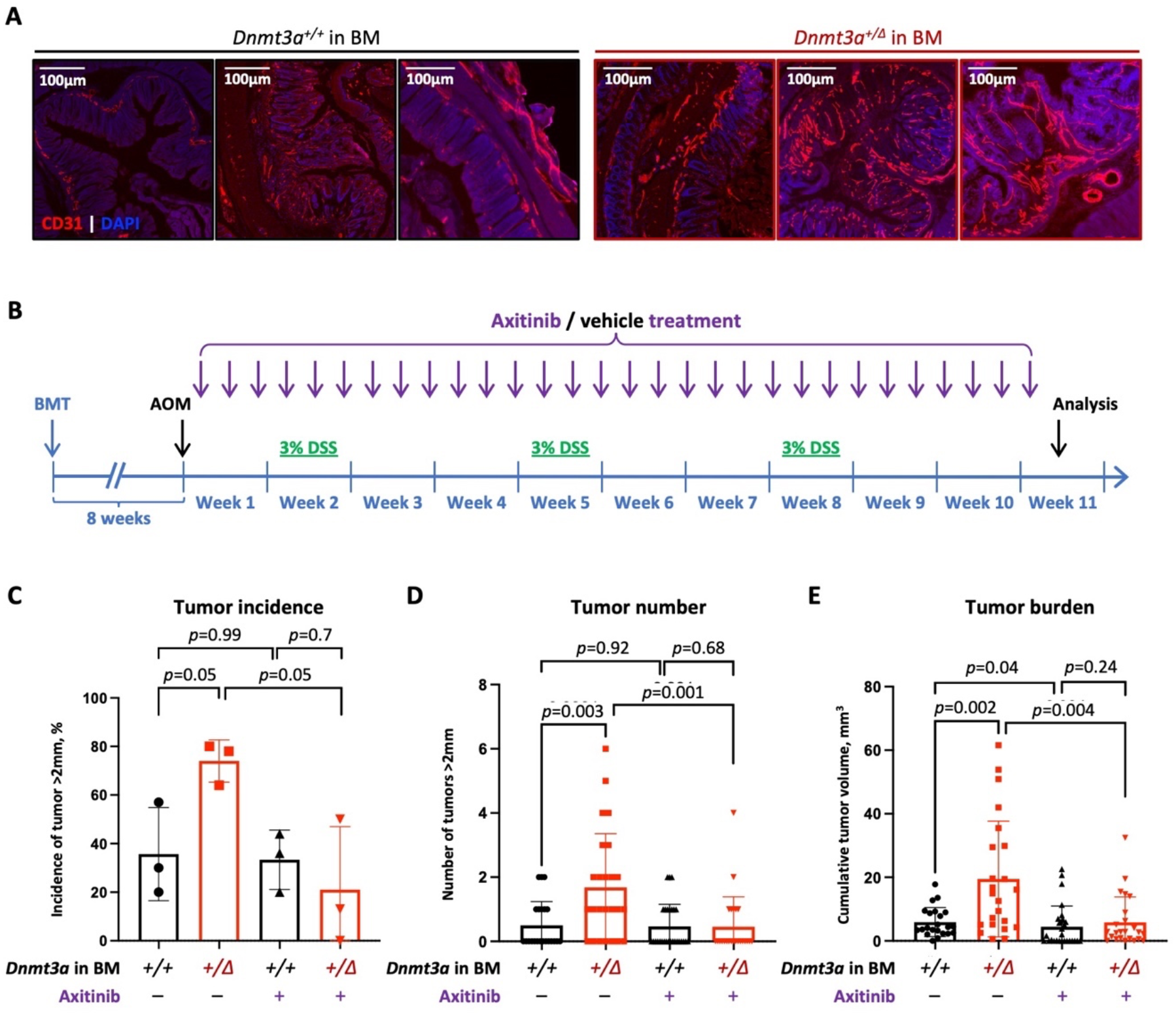
Targeting increased tumor angiogenesis in *Dnmt3a^+/Δ^*-CH animals mitigates accentuated CAC phenotype. (**A**) Representative examples of CAC tumors immunofluorescently stained for endothelial marker CD31 (red) demonstrate enhanced cancer angiogenesis in colons from *Dnmt3a^+/Δ^* chimerae compared to *Dnmt3a*^+/+^-transplanted controls; DAPI was used to visualize nuclei (blue). Bar – 100μm. (**B-E**) Treatment with small molecule angiogenesis inhibitor axitinib (25mg/kg per os, 3 times per week) mitigates elevated CAC tumorigenesis in *Dnmt3a^+/Δ^*-CH mice. Experimental timeline of axitinib treatment in CH-CAC animal model (**B**). Proportion of animals with large tumors (>2mm in diameter) after axitinib or vehicle control treatment in 3 independent experiments (**C**, Welch’s t test); number of large tumors (>2mm in diameter) per animal, with or without axitinib treatment (**D**, n=22-28, Welch’s *t* test, or Mann-Whitney test); and cumulative tumor burden per animal (**E**, n=22-28, Welch’s t test, or Mann-Whitney test).

## DISCUSSION

In patients with non-hematologic cancers, the incidence of CH is significantly higher than in age-matched tumor-free population (33,34), raising the clinically and biologically important question if these two conditions are causally linked. With approximately 10^5^ HSCs in adult bone marrow, an estimated one million of protein-coding mutations are acquired by the stem cell pool by the age of 60 (88–90). While most mutations are functionally neutral or occasionally detrimental, some genetic alterations may confer fitness advantage, allowing select HSCs to outcompete their peers and dominantly contribute to the pool of mature blood cells in the periphery (13,90–93). Such clonal expansion with an individual HSC at the apex and its normal mature progeny, termed clonal hematopoiesis (CH), is increasingly prevalent in advanced age individuals (11,12,57,91,94).

CH is commonly detected by presence of mutations in the bone marrow or peripheral blood, quantified by calculating variant allele frequency (VAF) (1–4). As most CH mutations were initially discovered in acute myeloid leukemia or other hematologic malignancies, it is not surprising that presence of CH may signify an increased risk of a future blood cancer (8,10,16,18–20,95). However, studies utilizing serial blood samples over multiple years of follow up suggest that this risk of leukemic progression is insufficiently high to explain increased overall mortality in individuals with CH (15,17). At the same time, genetic alterations in BM-derived cells are likely to affect their function and contribute to disease conditions outside of the hematopoietic system such as atherosclerosis, with experimental validation emerging (23,73–75). Yet in patients with solid tumors, presence of confounding factors that are known to contribute to both solid tumor and CH development (use of tobacco products, prior radiation or chemotherapy exposures including PARP inhibitors) further obscures the inference of causality in clinical trial data (18,34,69,70,96,97), necessitating experimental testing of the cause-effect relationship in model systems. These challenges were addressed in the current study.

The current study investigated the role of *DNMT3A*-driven clonal hematopoiesis in the aggressive phenotype of coincident colitis-associated colon cancer. We combined a well characterized mouse model of CAC with a BMT-based approach to experimental CH driven by heterozygous *Dnmt3a* loss. Azoxymethane/dextran sulfate sodium salt (AOM/DSS) CAC is a gold-standard autochthonous model with high penetrance and predictable latency (65–67). The use of immunocompetent animals preserves all modes of tumor-microenvironment interactions. AOM/DSS chemically-induced CAC produces a spectrum of mutations that captures genetic landscapes observed in human colon cancer (65) and recapitulates human pathology. This unique tool enables rigorous interrogation into CH involvement in the pathogenesis of solid tumors driven by unrelated genetic alterations, which can be readily extended to other cancer types. Our results demonstrate increased tumor number and burden, more severe histopathology and dysregulated signaling pathways in animals with experimental CH driven by heterozygous *Dnmt3a* loss. Although in our study using CAC model *Dnmt3a*-CH promoted both tumor initiation and progression, consistent with prior pan-cancer retrospective clinical observations (34), this effect is likely context specific. Thus, in patients with metastatic colorectal cancer enrolled in the FIRE-3 clinical trial, presence of CH, and CH-*DNMT3A* specifically, was associated with extended overall survival (98). Consistently, in our clinical cohort, patients with CH presented with more advanced disease at cancer diagnosis, although the study was not powered to evaluate survival. These contrasting observations may reflect disparate effects of *DNMT3A* loss in different branches of the immune system within the tumor microenvironment tipping the balance between anti-tumorigenic and pro-resolving phenotypes (46,99). While outside the scope of this study, these specific mechanisms remain to be investigated in the future. Further, the effects of *Dnmt3a*-CH likely extend beyond anti-cancer immunity, illustrated by the striking finding of enhanced tumor angiogenesis in animals with experimental CH. In support of this, targeted inhibition of angiogenesis with an FDA-approved small molecule tyrosine kinase inhibitor axitinib abrogated the CAC tumor-promoting effect of experimental *Dnmt3a*-CH. These results identify an actionable therapeutic strategy to mitigate the tumor-promoting effect of coincident *DNMT3A*-CH that can be tested in the clinical setting.

Together, these results highlight a significant difference of molecular pathophysiology effected by genetic alterations within the hematopoietic compartment, despite identical mode of CAC induction. It indicates that alteration of *Dnmt3a* in the bone marrow has a profound impact on molecular pathogenesis of CAC through a non-tumor-cell-autonomous mechanisms. These findings, for the first time, solidify the causal relationship between CH and the severity of solid tumors, and identify potential therapeutic strategies. Further study is needed to investigate underlying molecular mechanisms including immune involvement.

## Author contributions

YF, CJ, DA, and OAG designed research; YF, RCN, ANZ, KNH, CMB, DES, KIK, CG, ZJZ, KV, HLCR, and OAG performed research; TR, RLB, and AR analyzed RNA-seq data; RY, WKC, and RLL provided clinical data; DK assisted with imaging studies; YF and OAG analyzed data and wrote the manuscript with input from all co-authors.

## Acknowledgements

This work was supported by NIH awards R00CA178191 and R01DK121831 (OAG), R01AR079449 (DK), R01AI067846 (DA), the Thomas H. Maren Junior Investigator Fund (OAG), the Harry T. Mangurian, Jr. Foundation (OAG), and UFHCC Interdisciplinary Pilot Award (OAG, CJ, DA). OAG is supported by the Edward P. Evans Foundation. Next generation sequencing and flow cytometry analyses were performed at the UF Interdisciplinary Center for Biotechnology Research (ICBR) RRIDs SCR_019145, SCR_019152, SCR_019120, SCR_019119. The authors thank Lidia Kulemina, PhD for editorial assistance. This work was also supported by an NCI Cancer Center Support Grant to MSK (P30 CA08748).

## METHODS

### Patients

Patients with colitis-associated cancers were identified from a database of genomically annotated CAC cases maintained under MSK IRB protocols 15-297 and WA0143-14. All participating patients signed informed written consent for matched tumor and normal sequencing (MSK IRB 12-245), and next-generation sequencing was performed with the MSK-IMPACT assay (100). MSK-IMPACT is a hybridization capture-based next generation assay encompassing all exons of >340 genes. It is validated and approved for clinical use by New York State Department of Health Clinical Laboratory Evaluation Program. The sequencing test utilizes genomic DNA extracted from formalin-fixed paraffin-embedded (FFPE) tumor tissue as well as matched patient blood samples. DNA is sheared, and DNA fragments are captured using custom probes. MSK-IMPACT contains most of the commonly reported CH genes with the exception that earlier versions of the panel did not contain PPM1D or SRSF2.

### Mice

Animals were housed at the University of Florida Cancer & Genetics Research Complex specific pathogen-free animal facility; all animal studies were approved by the University of Florida Institutional Animal Care and Use Committee under protocol #201909474. A conditional *Dnmt3a* knock-out line was previously described (42,101). *Dnmt3a^+/fl^ and Dnmt3a^+/+^ mice* with *Mx1*-Cre deletor on a C57BL6/J background were generated *via* in house breeding. To achieve inducible hematopoietic-specific excision, *Dnmt3a^+/fl^:Mx1-Cre* mice received five intraperitoneal injections of poly(I:C) (InvivoGen, #tlrl-pic-5). *Mx1*-Cre-driven recombination was validated by PCR using genomic DNA from peripheral blood mononuclear cells. To model experimental CH driven by *Dnmt3a* LOF, BM cells from animals with either heterozygous loss of *Dnmt3a* (*Dnmt3a^+/Δ^*) or wild-type controls (*Dnmt3a*^+/+^) marked by CD45.2 were transplanted into lethally irradiated (10.5 Gy split dose) 6-week old congenic wild-type CD45.1 recipients (The Jackson Laboratory, strain #002014) through tail vein injection. Successful engraftment was confirmed by CD45.1/CD45.2 peripheral blood chimerism 2 months after BMT. Mice of different genotypes were co-housed to control for possible cage effects. Animals received unique IDs for the purpose of blinding; investigators were unblinded after analyses were complete. Peripheral blood was collected by submandibular puncture. Complete blood counts were obtained using HESKA HT5 automated veterinary hematology analyzer. Both male and female recipients were used with equivalent results.

### Colitis-associated colon cancer (CAC) induction

CAC is induced in a 10 week-long protocol. Successfully engrafted mice received a single dose of azoxymethane (AOM, Sigma-Aldrich, Cat# A4586, 10mg/kg) by intraperitoneal injection. One week later, Dextran Sodium Sulfate (DSS, Alfa Aesar, AAJ6360622, MW *ca* 40,000) was provided in sterile drinking water at 3% (w/v) as the sole source of water during each of the three DSS treatment cycles with two weeks of sterile drinking water between cycles. As the effect of DSS may vary between mouse strains, genders, and housing facilities, each lot was tested to determine the optimal DSS dosing. Body weights were monitored daily during DSS administration, animals losing more than 25% body weight were euthanized as past humane endpoint approved by the IACUC. Colon microbiota was normalized by mixing cage bedding of all experimental groups every other week to eliminate cage-to-cage variability. Since AOM is a systemic carcinogen, all animals were bled monthly to monitor for hematologic malignancies. Animals with hematopoietic disorders diagnosed by CBCs and flow cytometry analysis of peripheral blood mature lineages (102) were excluded as potential confounders. At day 70, animals were bled and euthanized for comprehensive analysis. Flow cytometry data were collected on a 5-laser 16-parameter BD LSR Fortessa instrument and analyzed by FlowJo v10, using the following antibody cocktail:

| MARKER | FLUOROPHORE | PER 100μL | CLONE | MANUFACTURE | CATALOG# |
| --- | --- | --- | --- | --- | --- |
| CD45.1 | PacBlue | 0.5 | A20 | BioLegend | 110722 |
| CD45.2 | APC | 0.3 | 104 | BioLegend | 109814 |
| B220 (CD45R) | FITC | 0.3 | RA3-6B2 | BioLegend | 103206 |
| CD4 | FITC | 0.5 | GK1.5 | BioLegend | 100406 |
| CD8 | FITC | 0.5 | 53-6.7 | BioLegend | 100706 |
| CD3 | FITC | 0.3 | 17A2 | BioLegend | 100706 |
| c-Kit (CD117) | PE | 0.5 | 2B8 | BioLegend | 105808 |
| B220 (CD45R) | APC-Cy7 | 0.3 | RA3-6B2 | BioLegend | 103224 |
| CD11b | APC-Cy7 | 0.1 | M1/70 | BioLegend | 101226 |

### Colonoscopy in live animals

Colonoscopy was performed after second and third DSS cycles using the Tele Pack Vet X mini-endoscopic system with rigid telescopes (KARL STORZ Veterinary Endoscopy, CA) as previously described as previously described (103–106). Briefly, mice were anesthetized with isoflurane and placed ventral side up on a heating pad. Endoscope was carefully inserted into the rectum up to 4cm under visual control with slow air flow to keep the colon inflated, and withdrawn slowly while recording the localization and size of colon abnormalities. Colonoscopy videos were independently scored by two blinded investigators using modified murine endoscopic index of CAC severity (MEICS) (78) that combines five scoring criteria (colonic wall thickening (0-3), vascular pattern (0-3), fibrin formation (0-3), stool consistency (0-3), and tumor diameter (0-5), for a total score ranging 0-17).

### Tissue dissection and processing

Colons were dissected immediately after euthanasia. After washing out the feces by 2-3ml sterile PBS, colons were laid on wet Whatman blotting paper and cut longitudinally. Tumors were counted and dimensions in millimeters measured with digital calipers used to calculate tumor volume V = (Width^2^ × Length)/2. For histological analysis, each dissected colon was swiss-rolled from the distal end.

### Immunohistochemistry

Swiss-rolled colons were fixed in 4% paraformaldehyde (PFA) in sterile PBS for 24 hours, transferred to 70% ethanol and stored at 4°C. Paraffin embedding, sectioning, and hematoxylin and eosin (H&E) staining, and Ki67 immunohistochemistry were performed at UF Molecular Pathology Core. Briefly, 4 μm sections were de-paraffinized, and treated by Trilogy (CELL MARQUE REF:920P-06) in a 95°C water bath for 25 minutes. Background Sniper (Biocare Medical #BS966M) was applied for 15 minutes to reduce background. Sections were incubated with rat anti-mouse Ki67 (1:50, DAKO #M7249) for 60 minutes, followed by NB Rabbit anti Rat 1:200 for 30 minutes. Stain was visualized using Mach 2 Rabbit HRP polymer (Biocare Medical # RHRP520L), and the DAB chromagen (Vector Laboratories, #SK-4105) with CAT hematoxylin counterstain (Biocare Medical #CATHE-M). Slides were scanned using Keyence BZ-X800 microscope and VHX series software.

### Immunofluorescent staining

Swiss-rolled colons were fixed as described above, placed in series of 10%-20%-30% sucrose solutions in PBS until sinking, and embedded in OCT compound (Fisher, Cat# 23730571). Frozen samples were cut into 12 μm thick sections at −20°C, bound to positively charged microscope slides (ASI, Cat# SM2575) at room temperature (RT), and stored at -20°C. For immunofluorescent staining, slides were thawed at RT for 20-30 min and tissue permeabilized with 0.2% Triton-X 100 in PBS at RT for 10 min, then thoroughly washed with PBS. After blocking with 10% donkey serum in PBS for 1 hour, slides were incubated with goat anti-mouse CD31 primary antibodies (R&D Systems, #AF3628; 1:100) in blocking buffer overnight in 4°C and stained with secondary AlexaFluor633 donkey anti-goat antibodies (Invitrogen, #A21082; 1:200) for 1 hour at RT, three washes with PBS after each incubation, and mounted with Prolong Gold Antifade with 4’,6-diamidino-2-phenylindole (DAPI) (Invitrogen, P36935) as a counterstain for nuclei. Images were acquired with a Leica DMi8 confocal microscope equipped with a DFC7000 camera using a 20× HC PL Fluotar objective (Leica). The LAS Navigator function was used to generate a merged image of the whole cross-section. All images were processed identically using Fiji/ImageJ (NIH).

### Sample preparation for RNA extraction, sequencing, and analysis

After dissection, each colon tumor was submerged in 200μl RNAlater (Thermo Scientific, AM7021) in 1.5ml Eppendorf tube, flash frozen in liquid nitrogen, and stored at −80°C. Tumor tissue was thawed on ice and transferred into a blaster tube containing 500μl buffer RLTplus (Quiagen,1053393) and 200μl glass beads (VWR, 12621-148), then homogenized using BeadBlaster 24 Microtube Homogenizer (Benchmark) at 6M/s for 30s three times, with 2 min breaks for cooling. Total cellular RNA was isolated with RNeasy Microprep kit (QIAGEN, 74004) and QC’d on a 4150 TapeStation (Agilent). RNA samples with RNA integrity number (RIN) 7.0 or higher were picked for library preparation and sequencing. RNA was subjected to standard Illumina-based RNAseq library preparation and sequenced on an Illumina NovaSeq6000 using a paired-end 100bp chemistry to an average depth of 49M reads/sample. Input sequences were trimmed with Trimmomatic; quality control was performed before and after trimming using FastQC. Retained reads (>96%) were aligned to mouse reference transcriptome mm10/build 38 using STAR 2.7.3a; gene and transcript expression (raw FPKM) were quantified using RSEM v1.2.31. From a transcript count matrix, differential gene expression was evaluated with DESeq2 using a log2FC cutoff of +/− 1 and a false discovery rate of 5% (107). Gene set enrichment analysis was conducted using the *fgsea* package in R. The volcano plot (Fig. 3A) and heatmap (Fig. 3C) were produced using the *EnhancedVolcano* and *pheatmap* R packages, respectively. All R code used to analyze RNA-sequencing data will be made publicly available at https://github.com/RobinsonTroy/CH_CAC_RNAseq.

### Data availability

Raw and processed RNA-seq data are deposited at GEO (GSE213178).

### Statistical analysis

Sample size calculation was based on the primary endpoint: tumor burden (mm^3^). The number of animals was chosen to ensure 90% power with 5% alpha to detect a difference between groups of one standard deviation (SD) or larger based on variability and technical drop-out rate observed in pilot experiments. All grouped data are presented as mean ± SD. Statistical significance was determined by unpaired parametric Student’s *t*-test and by non-parametric Mann-Whitney test after testing for normal distribution. For samples with significantly different variances, Welch’s correction was applied. Statistical analyses and visualization of the data were performed using Prism 9.0.2 (GraphPad Inc.). For pair-wise comparisons, *p* values equal or less than 0.05 were considered significant.

### Rigor and reproducibility

For the entire study, both female and male animals were used to control for gender-specific effects. All experimental animals from different genotype groups were co-housed to mitigate potential cage effects. Other variables were kept consistent in all cages and experiments, such as microbiota and DSS quality. Animals developing hematologic malignancies were excluded from analysis as potential confounders. Blinding was achieved by assigning random codes to each animal and sample prior to analysis; investigators were unblinded after results had been recorded.

## Supplementary material

Supplementary tables S1, S2, and S3 summarize RNA-seq data from colon tumors derived from 3 mice with *Dnmt3a*-CH and 4 WT-engrafted control animals, including normalized gene counts (Suppl. Table S1), differentially expressed genes (Suppl. Table S2), and enriched gene sets (Suppl. Table S3).

**SUPPLEMENTARY MATERIAL** (available as separate files online)

**Supplementary Table S1. Normalized gene counts in colon tumors from *Dnmt3a^+/Δ^*-CH mice (*n*=4) and *Dnmt3a*^+/+^-grafted controls (*n*=3)** (RNA-seq)

**Supplementary Table S2. Differentially expressed genes in colon tumors from *Dnmt3a^+/Δ^*-CH mice *vs Dnmt3a*^+/+^-grafted controls** (fold change >2, *p*-adjusted <0.05)

**Supplementary Table S3. HALLMARK pathways (MSigDB) enriched among differentially expressed genes between colon tumors from *Dnmt3a^+/Δ^*-CH and *Dnmt3a*^+/+^-grafted control mice**

## Notes

**Conflict of interest:** RLB: speakers bureau, MissionBio, unrelated to this work. RLL is on the supervisory board of Qiagen and is a scientific advisor to Imago, Mission Bio, Zentalis, Ajax, Auron, Prelude, C4 Therapeutics, and Isoplexis. He receives research support from and consulted for Celgene and Roche and has consulted for Incyte, Janssen, Astra Zeneca, Morphosys, and Novartis. He has received honoraria from Astra Zeneca, Roche, Lilly, and Amgen for invited lectures and from Gilead for grant reviews. All other authors declare no financial conflicts of interest.

### Competing Interest Statement

RLB: speakers bureau, MissionBio, unrelated to this work. RLL is on the supervisory board of Qiagen and is a scientific advisor to Imago, Mission Bio, Zentalis, Ajax, Auron, Prelude, C4 Therapeutics, and Isoplexis. He receives research support from and consulted for Celgene and Roche and has consulted for Incyte, Janssen, Astra Zeneca, Morphosys, and Novartis. He has received honoraria from Astra Zeneca, Roche, Lilly, and Amgen for invited lectures and from Gilead for grant reviews. All other authors declare no financial conflicts of interest.

